# Retrospective Analysis of Mobile Colistin Resistance and Plasmid-Mediated Quinolone Resistance Genes in *Escherichia coli*: China, 1993-2019

**DOI:** 10.1101/2023.10.02.560564

**Authors:** Yujin Wang, Dewei Sun, Zhengzhong Xu, Xin’an Jiao, Xiang Chen

**Author notes:** Author order was determined by drawing straws.

## Abstract

Plasmid-mediated quinolone resistance (PMQR) genes and mobile colistin resistance (MCR) genes in *Escherichia coli* (*E. coli*) have shown an increasing prevalence in recent years. This rise is particularly notable in China and several Southeast Asian countries, representing a significant threat to public health. In the present study, we conducted a retrospective analysis of MCR genes (*mcr-1*, *mcr-2*, *mcr-3*, *mcr-4*, and *mcr-5*) and PMQR genes (*qnrA*, *qnrB*, *qnrC*, *qnrD*, *qnrE1*, *qnrVC*, *qnrS*, *aac(6’)-Ib-cr*, *qepA*, and *oqxAB*). From the 3,663 *E. coli* isolates examined, 1,613 (44.0%) tested positive for PMQR genes, either individually or in combination. Meanwhile, 262 isolates (7.0%) carried MCR genes. Minimum inhibitory concentration (MIC) analyses of 17 antibiotics for the MCR gene-carrying strains revealed universal multidrug resistance. Resistance to polymyxin varied between 4 μg/mL and 64 μg/mL, with MIC50 and MIC90 values at 8 μg/mL and 16 μg/mL, respectively. Additionally, fluctuations in the detection rates of these resistant genes correlated with the introduction of antibiotic policies, host origin, temporal trends, and geographical distribution.

**IMPORTANCE:** Antimicrobial resistance (AMR) has emerged as a threat to global health and development, and *E. coli* resistance remains an ongoing clinical challenge. However, the efficacy of colistin and quinolones has been compromised by the presence of various MCR and PMQR genes. To elucidate the prevalence and distribution of MCR and PMQR genes in *E. coli* isolates, we analyzed 3,663 *E. coli* isolates from 29 provinces in China from 1993 to 2019. The results show that 1,613 (44.0%) tested positive for PMQR genes, 262 isolates (7.0%) carried MCR genes. And MICs analyses of 17 antibiotics for the MCR gene-carrying strains revealed universal multidrug resistance. These indicated prohibiting or strictly curtailing antimicrobial use in animal or human is urgently needed to reduce the growing threat from AMR.

## INTRODUCTION

The discovery and application of antibiotics mark a monumental shift in global health, revolutionizing modern medicine, agriculture, and animal husbandry [1]. Yet, the extensive use of these antibiotics has hastened the emergence of bacterial resistance, giving rise to multi-resistant bacteria [2]. The World Health Organization (WHO) has identified antimicrobial resistance as an urgent global public health crisis that demands immediate attention [3].

*E. coli* is a Gram-negative bacterium commonly found as part of the normal intestinal flora, but it can also cause a range of diseases [4]. While *E. coli* is naturally sensitive to most clinical antimicrobials, it possesses a significant ability to acquire resistance genes. These genes are often associated with mobile genetic elements like plasmids and transposons, enabling their transfer between bacteria from different phylogenetic lineages. Consequently, *E. coli* drug resistance remains a persistent clinical challenge [5,6]. In 2021, Gram-negative bacteria comprised 71.1% of all drug-resistant bacterial detections in China, with *E. coli* topping the list, accounting for 29.2% of all isolates [10]. Furthermore, the WHO ranks it third in statistics for antibiotic-resistant bacteria, at a concerning 76.5% [7].

Polymyxin is a polypeptide antibiotic derived from the culture solution of *Bacillus polymyxa var*. It encompasses five distinct members: polymyxin A, B, C, D, and E. Among these, polymyxin B and polymyxin E are most commonly utilized in clinical settings. Given that many multidrug-resistant Gram-negative bacteria remain susceptible to polymyxins, these antibiotics are often regarded as the “last line of defense” in treating such resistant strains [8]. Resistance of Gram-negative bacteria to polymyxin can be categorized into innate resistance and acquired resistance [9].

Breaking from earlier understood mechanisms of polymyxin resistance, Liu et al. are the first to report that the *mcr-1* gene can disseminate polymyxin resistance via plasmids, drawing significant attention [10,11]. Since then, 10 mobile colistin resistance (MCR)-family genes (*mcr-1* to *mcr-10*) and their variants have been identified. Notably, while *mcr-6* is chromosome-located, the remaining MCR genes are found on plasmids, facilitating their horizontal transfer [12,13]. This has enabled their global spread, causing a wide range of bacteria to exhibit increased tolerance to colistin.

Fluoroquinolones (FQs) stand out as some of the most successful antibiotics in recent history. With over three decades of clinical use, they have become indispensable in the repertoire of clinical treatments. However, resistance to FQs is on the rise [14]. *qnr* is the first plasmid-mediated quinolone resistance (PMQR) gene identified [15]. Currently, around 100 *qnr* variants have been cataloged, grouped into six distinct families: *qnrA*, *qnrB*, *qnrS*, *qnrC*, *qnrD*, and *qnrVC*. In 2006, another PMQR mechanism was discovered, the most notable being *aac(6’)-Ib-cr*. This gene diminishes the efficacy of ciprofloxacin and norfloxacin by modifying them [16]. Subsequently, plasmid-encoded efflux pumps, *qepA* and *oqxAB*, are identified [17]. Pathogens harboring PMQR genes are more prone to genetic mutations, which can confer high levels of antibiotic resistance. Moreover, the co-presence of PMQR genes alongside extended-spectrum β-lactamase (ESBL) genes may restrict treatment alternatives for infections by ESBL-producing bacteria [18].

Over recent decades, the proliferation of antibiotic resistance in *Enterobacteriaceae* has posed a formidable challenge to medical treatment. As one of the global leaders in both antibiotic production and consumption, China is at the forefront of this crisis, making it imperative to thoroughly investigate the prevalence of antibiotic-resistant genes within the nation [19].

In this retrospective study, we analyzed 3,663 *E. coli* strains collected from 31 provinces (including municipalities and autonomous regions) spanning 1993 to 2019. We specifically detected the presence of PMQR genes (*qnrA*, *qnrB*, *qnrC*, *qnrD*, *qnrE1*, *qnrVC*, *qnrS*, *aac(6)-Ib-cr*, *qepA*, and *oqxAB*) and MCR genes (*mcr-1*, *mcr-2*, *mcr-3*, *mcr-4*, and *mcr-5*). Through polymerase chain reaction (PCR) detection and multidimensional analysis, we elucidated the epidemiological characteristics of MCR and PMQR genes in *E. coli*, deduced their primary patterns of spread, and compiled comprehensive epidemiological data. This information was crucial in curbing the further dissemination of MCR and PMQR genes, enhancing the mitigation of drug-resistant strains, and safeguarding public health.

## RESULTS

### Overview of MCR and PMQR genes in E. coli during 1993-2019

Of the 3,663 *E. coli* isolates collected from various sources between 1993 and 2019, 262 strains (7.2%) contained *mcr* genes: 258 strains had *mcr-1* (7.0%), five strains had *mcr-3* (0.1%), and one strain possessed both *mcr-1* and *mcr-3*. Moreover, 1,613 strains (44.0%) tested positive for PMQR genes. Specifically, the presence of *qnrA* (0.3%), *qnrB* (3.8%), *qnrS* (17.1%), *qnrVC* (0.4%), *aac(6’)-Ib-cr* (7.3%), *qepA* (0.9%), and *oqxAB* (21.9%) was identified either individually or in combination (Table 1 and Supplemental Table S3). None of the strains contained *qnrC*, *qnrD*, *qnrE1*, *mcr-2*, *mcr-4*, or *mcr-5*. Additionally, a polygenic distribution was observed that 9.1% (332/3,663) of isolates contained multiple PMQR genes concurrently (Supplemental Table S4). Besides, 7.8% (287/3,663) had two PMQR genes, and 1.2% (45/3,663) contained three PMQR genes. The combinations *oqxAB* + *qnrS* (3.2%) and *oqxAB* + *aac(6’)-Ib-cr* (2.4%) were prevalent, followed by *qnrS* + *aac(6’)-Ib-cr* (0.9%). Among strains with three PMQR genes, the combination *oqxAB* + *qnrS* + *aac(6’)-Ib-cr* was the most common (0.9%).

**Table 1.** The overall resistance of MCR-detection strains.

| Antibiotics | MIC value (µg/mL) |  |  |  | Resistance rate |
| --- | --- | --- | --- | --- | --- |
|  | Range |  | MIC<sub>50</sub> | MIC<sub>90</sub> |  |
|  | Min | Max |  |  |  |
| Colistin | 4 | >64 | 8 | 16 | 100% (262) |
| Ampicillin | 4 | >128 | >128 | >128 | 95.0% (249) |
| CTX | 0.03 | >128 | 16 | >128 | 73.3% (192) |
| Cefazolin | 0.5 | >128 | 128 | >128 | 78.2% (205) |
| Aztreonam | 0.06 | >128 | 32 | >128 | 70.6% (185) |
| Meropenem | 0.015 | 16 | 0.0625 | 0.125 | 2.3% (6) |
| Gentamicin | 0.125 | >128 | 16 | >128 | 56.9% (149) |
| Streptomycin | 1 | >256 | 126 | >256 | 63.7% (167) |
| Amikacin | 0.25 | >256 | 2 | >256 | 22.9% (60) |
| Tetracycline | 1 | >128 | 128 | >128 | 94.3% (247) |
| SXT | 1 | >32 | >32 | >32 | 97.7% (256) |
| Fosfomycin | 4 | >512 | >512 | >512 | 68.7% (180) |
| Ciprofloxacin | 0.008 | >64 | 32 | >64 | 82.4% (216) |
| Nalidixic acid | 2 | >256 | 256 | >256 | 93.1% (244) |
| Chloramphenicol | 2 | >128 | >128 | >128 | 85.1% (223) |
| Florfenicol | 1 | >128 | >128 | >128 | 77.1% (202) |
| Nitrofurantoin | 4 | 256 | 64 | 128 | 18.3% (48) |

Among the MCR-positive strains, 142 isolates (3.9%) also harbored PMQR genes (Table S4). For instance, 1.8% (67/3,663) contained *mcr-1* + *oqxAB*, and 0.9% (34/3,663) had *mcr-1* + *oqxAB* + *qnrS*. A particular strain sourced from pigs carried four resistance genes: *mcr-1*, *qnrS*, *oqxAB*, and *aac(6’)-Ib-cr*. Moreover, one strain possessed both *mcr-3* and *qnrS*, while another had *mcr-1*, *mcr-3*, and *qnrS*.

### Prevalence of PMQR and MCR genes in different hosts

The prevalence of *mcr* in 3,663 strains of *E. coli* from various host sources is detailed in (Table S5). The detection rate of *mcr-1* in pig isolates was the highest at 11.5% (97/846). This was significantly higher than any other host. It was followed by goose isolates at 8.0% (20/250) and chicken isolates at 7.5% (115/1532). The prevalence of MCR genes in poultry isolates was 7.5% (138/1,849) and 9.9% (104/1,049) in livestock isolates. Out of the human samples, 1.5% (9/614) tested positive for MCR genes, and among monkey samples, one strain or 1.1% (1/92) was *mcr-1*-positive.

The detection rate of PMQR genes in pig isolates was notably high at 76.4% (647/846), succeeded by goose isolates at 66.8% (167/250) and sheep isolates at 55.5% (50/90). The PMQR genes were identified in 52.0% (961/1,849) of poultry samples, 71.4% (749/1049) of livestock samples, and 22.3% (137/614) of human samples. The genes *oqxAB* and *qnrS* were detected as early as 1994 and 2003 in chickens or pigs, with detection rates higher than other genes at 21.9% and 17.1%, respectively. The earliest detection years for *mcr-1* and *mcr-3* were 2009 and 2016, respectively. Additionally, a new PMQR gene, *qnrVC4*, was first identified in *E. coli* in 2017 from a chicken (two samples) and a pig (one sample). Its detection rate stood at 0.3% (3/820) (Table S6).

### Geographical distribution of PMQR and MCR genes in China

In this study, the prevalence of *mcr-1* exceeded 15% in Central China, followed by Northeast China (above 10%), and then by East and South China (Fig. 1). However, North China, Southwest China, and Northwest China displayed lower prevalence rates. When regions were segmented according to the “Heihe-Tengchong line”, the *mcr-1*-positive isolates were significantly more frequent in the eastern part than in the western region. A total of five strains tested positive for *mcr-3* across three provinces: Jiangsu (two strains), Henan (two strains), and Qinghai (one strain) (Fig. 2).

**Figure 1.**
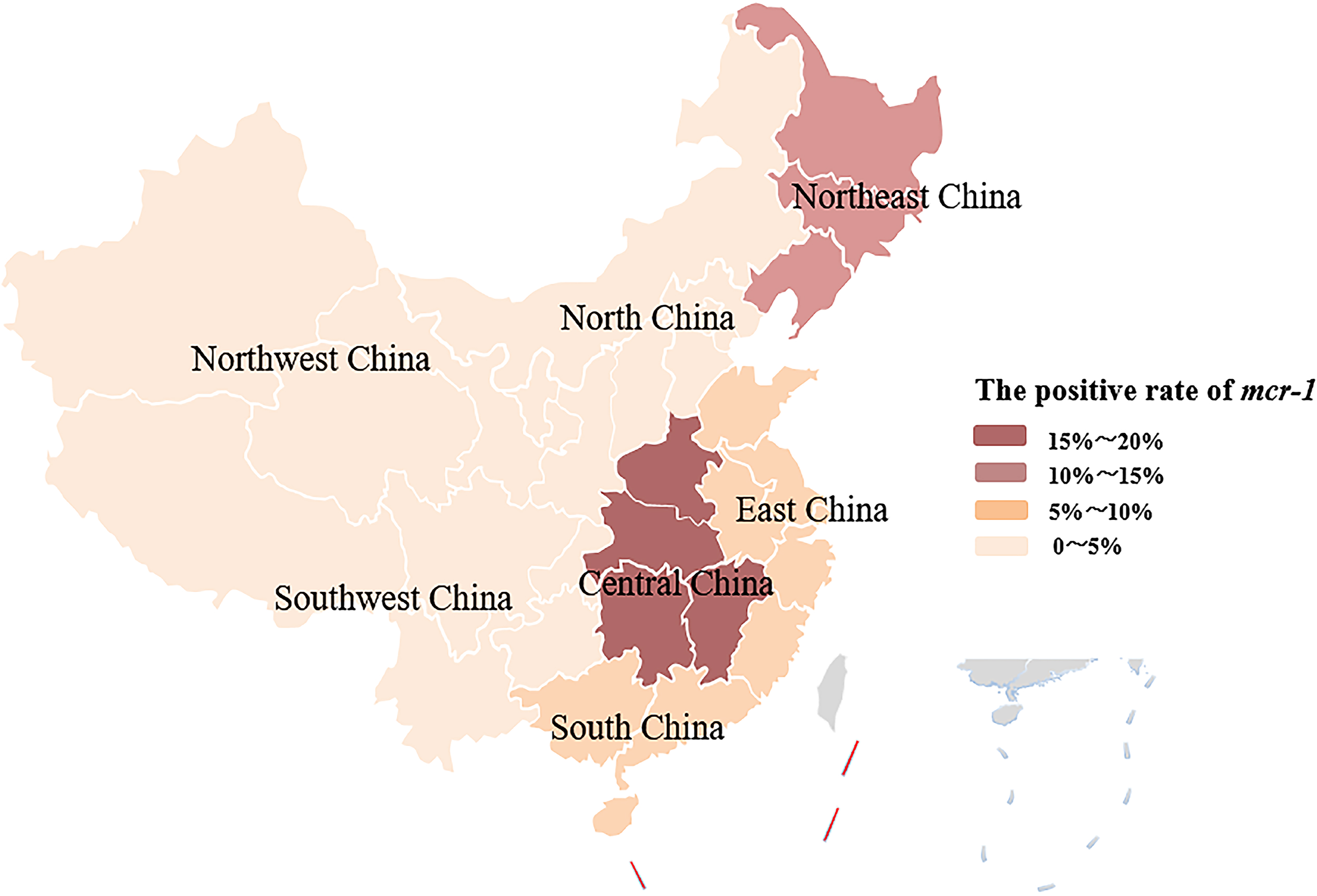
The prevalence and distribution of *mcr-1* in China from 1993 to 2018.

**Figure 2.**
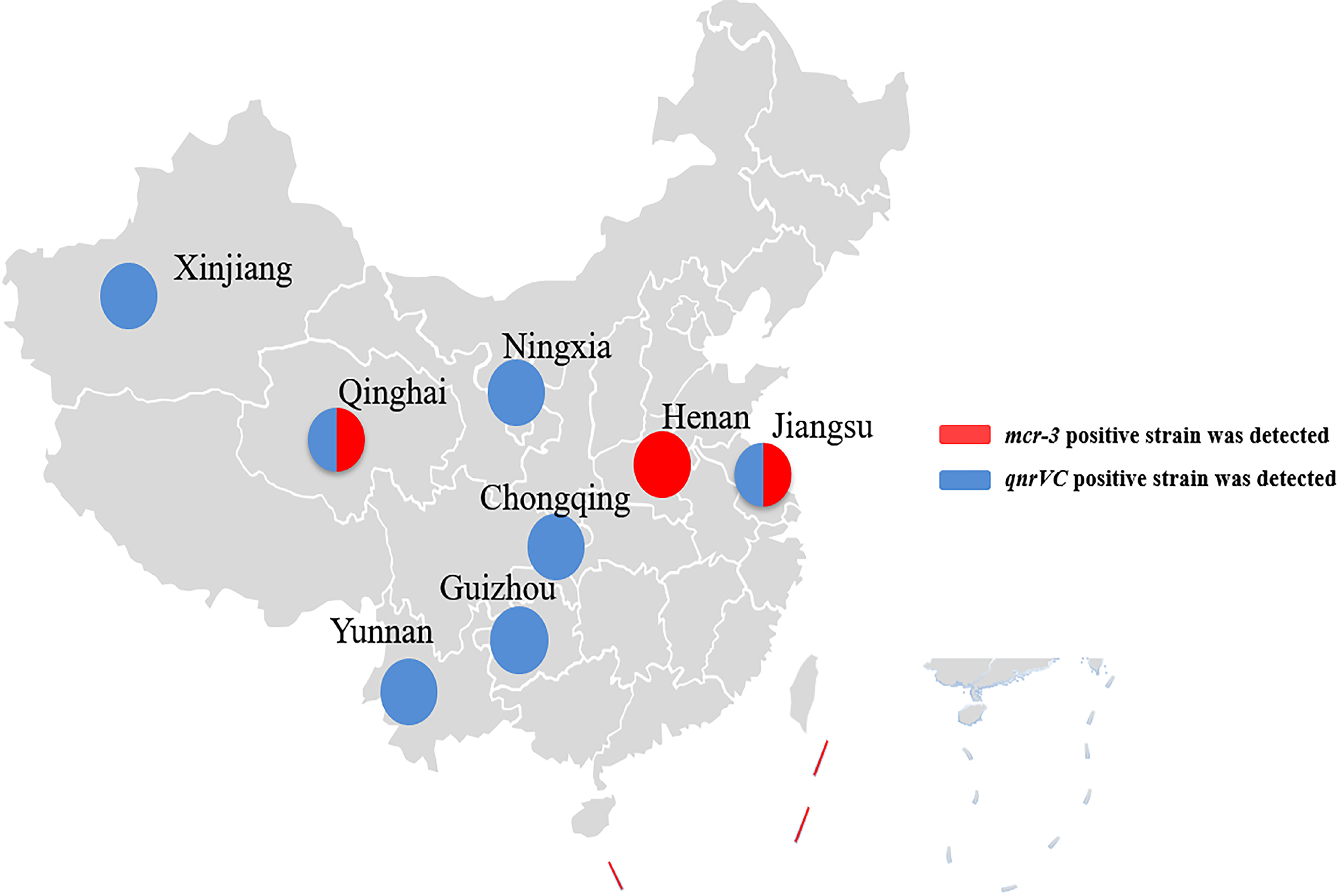
The prevalence of novel plasmid-mediated resistance gene *mcr-3* and *qnrVC* in China.

The overall detection rate of PMQR genes was highest in Northeast China, surpassing 20%. This was closely followed by Central China and Southwest China, both registering over 15% (Fig. 1). The *aac(6’)-Ib-cr* detection rate peaked in Central China. The novel PMQR gene, *qnrVC*, was identified in seven provinces: Xinjiang (two strains), Ningxia (one strain), Yunnan (four strains), Qinghai (two strains), Guizhou (one strain), Chongqing (one strain), and Jiangsu (five strains) (Fig. 3).

**Figure 3.**
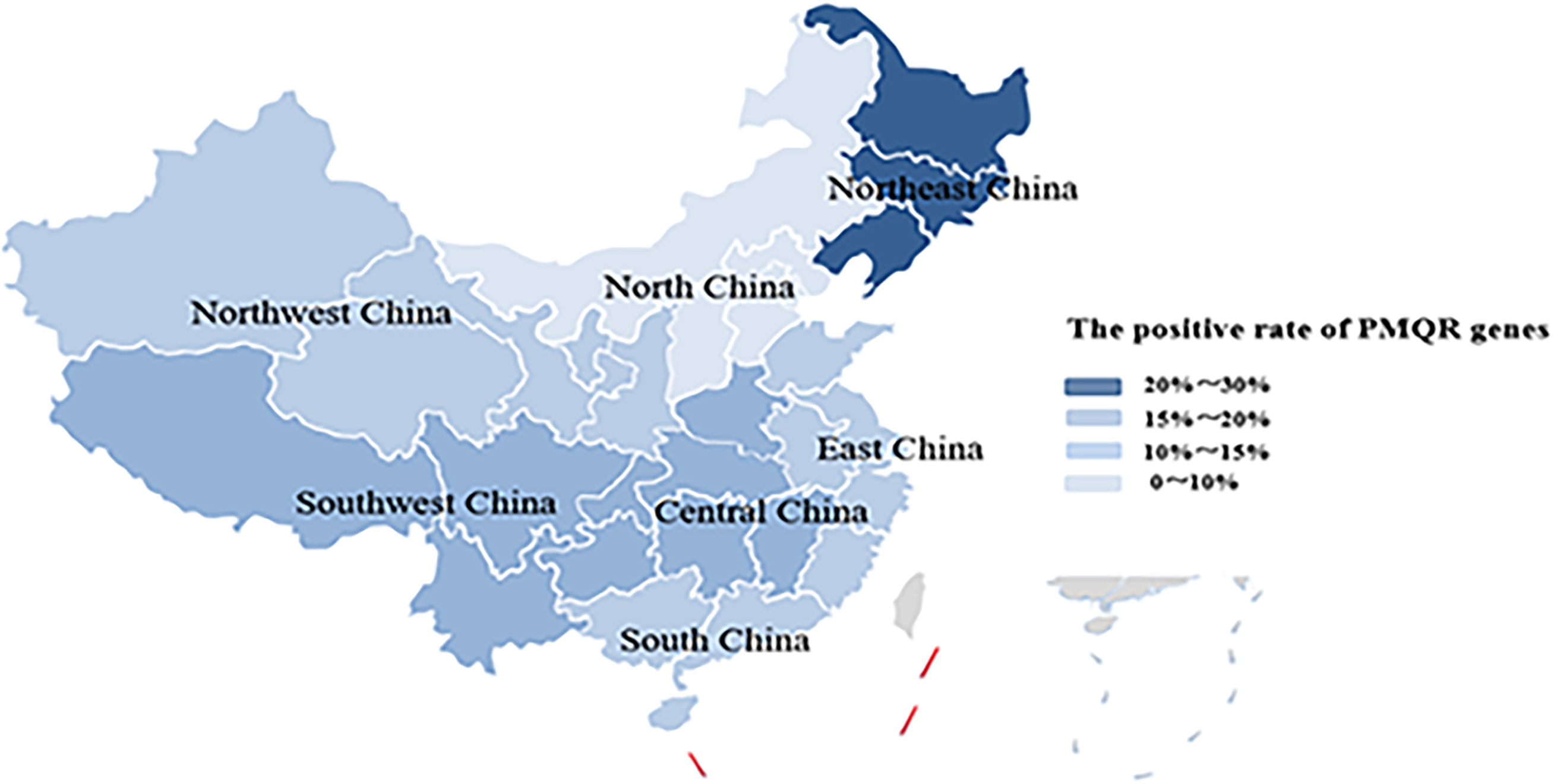
The prevalence and distribution of PMQR genes in China from 1993 to 2018.

### Temporal changes of PMQR and MCR genes

When analyzing the relationship between the number of veterinary drug approvals and the prevalence of *mcr-1* over time (Fig. 4), a distinct correlation emerged. The rate of *mcr-1* positivity appeared to correspond closely with the number of veterinary drugs sanctioned in any given year. This finding suggested that the increased approval of veterinary drugs might be influencing the prevalence of the *mcr-1* gene. *mcr-1* was first identified in chicken strains in 2009. Following this, the detection rate of *mcr-1* surged from 2009 to 2013, peaking at 16.8% (34/202) in 2013. Between 2014 and 2015, this rate consistently hovered around 16%. However, after the 2017 ban on using colistin as an animal growth promoter (Fig. S1 at http://www.ivdc.org.cn/), the prevalence of *mcr-1* swiftly declined to 5.0% (17/339) by 2019 [23].

**Figure 4.**
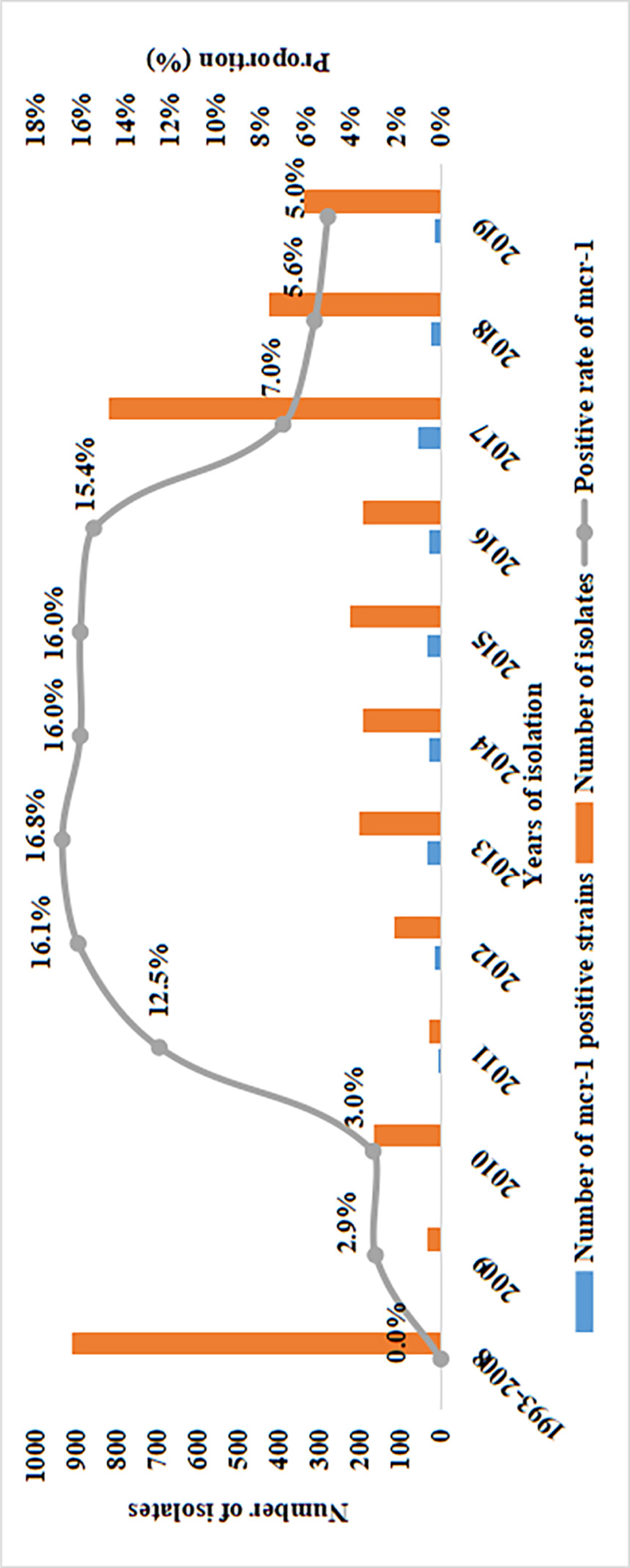
The detection rate of *mcr-1* in *E. coli* between 1993 and 2019.

Historical data suggested that the overall detection rate of *E. coli* in samples was high from 1993 to 2000. Even though the detection rate of *E. coli* in samples significantly dropped after the approval of enrofloxacin in the late 1980s, the carrying rates of *qnr* and *oqxAB* began to climb. By 2013, *oqxAB*-positive strains reached their zenith at 45.5% (92/202) (Fig. 5). In response to such trends, China has been progressively curtailing the approval of veterinary drugs (Fig. S2 at http://www.ivdc.org.cn/). In subsequent years, the detection rate of *oqxAB* consistently fell. By 2017, the rate dwindled to just 13.3% (109/820). Subsequently, three types of veterinary drugs were prohibited for use in food animals, quinolinol being one of them (details available at http://www.moa.gov.cn/nybgb/2015/jiuqi/201712/t20171219_6103873.htm). Contrary to expectations, the prevalence of *oqxAB* and *qnr*, particularly *qnr*, surged (Figure 6). In 2018, the detection rate for *qnrS* was a substantial 35.1% (150/427). The *qnrB* rate was minimal prior to 2016 but exhibited a modest uptick in 2017 and 2018, recording 8.4% and 9.8%, respectively.

**Figure 5.**
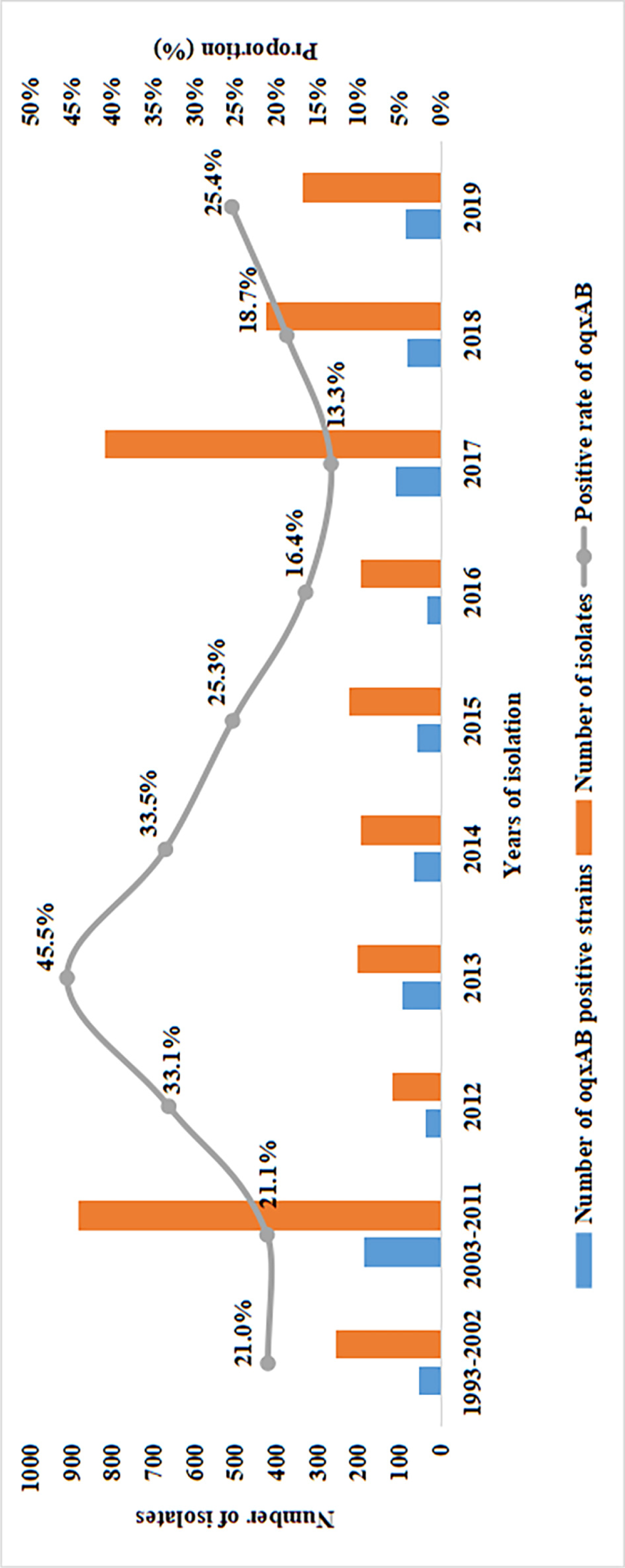
The detection rate of *oqxAB* in *E. coli* between 1993 and 2019.

**Figure 6.**
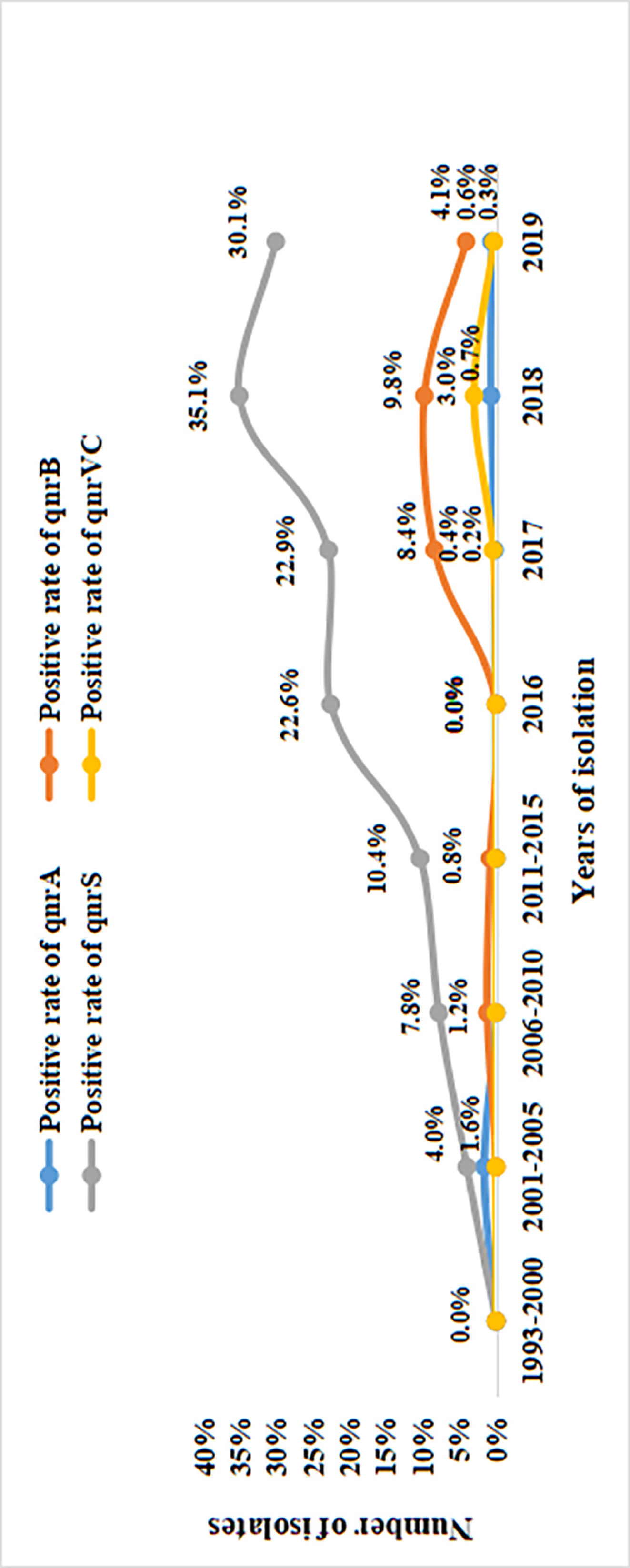
The detection rate of *qnr* genes in *E. coli* between 1993 and 2019.

### Antimicrobial susceptibility of MCR gene isolates

We assessed the MICs of 17 antimicrobial drugs against 262 MCR-positive isolates. All of these isolates displayed multidrug resistance (Table 1). Each MCR-positive strain demonstrated resistance to polymyxin, resulting in a 100% match rate between MIC phenotype and genotype. Resistance values ranged from 4 μg/mL to >64 μg/mL. The MIC_50_ and MIC_90_ values were 8 μg/mL and 16 μg/mL, respectively. For six strains, the MIC of polymyxin exceeded 64 μg/mL. In the context of β-lactam drugs, the resistance rates were as follows: ampicillin at 95%, cefotaxime at 73.3%, cefazolin at 78.2%, and aztreonam at 70.6%. Notably, six MCR-positive strains were simultaneously resistant to meropenem, with a resistance rate of 2.3% and the highest resistance value being 16 μg/mL. Regarding aminoglycosides, resistance rates were 56.9% for gentamicin, 63.7% for streptomycin, and 22.9% for amikacin. For quinolones, ciprofloxacin had a resistance rate of 82.4%, while nalidixic acid stood at 93.1%. Furthermore, resistance rates for other drugs were as follows: tetracycline at 94.3%, cotrimoxazole at 97.7%, fosfomycin at 68.7%, chloramphenicol at 85.1%, florfenicol at 77.1%, and nitrofurantoin at 18.3%.

From an overall drug resistance standpoint, isolates with MCR genes exhibited multidrug resistance. The resistance rate for most drugs hovered around 50% and often exceeded 80%. Only three drugs, meropenem, amikacin, and nitrofurantoin, demonstrated commendable antibacterial effects with resistance rates below 30%. Meropenem stood out with a notably low resistance rate of just 2.3%.

When analyzing the 262 MCR-positive strains based on host differences, our findings indicated (Table 2) that human MCR-positive strains had significantly lower drug resistance rates compared to other hosts. Whether considering β-lactam, aminoglycosides, or fosfomycin, chicken strains consistently displayed higher resistance rates than those from pigs. This was particularly evident for amikacin (34.5% vs. 11%) and fosfomycin (80.2% vs. 61%).

**Table 2.** Multidrug resistance of MCR-positive strains from different hosts.

| Resistance rate | Chickens<br>(n=116) | Ducks &<br>Geese<br>(n=23) | Pigs<br>(n=100) | Cattle &<br>Goat (n=7) | Humans<br>(n=9) | Other<br>animals<br>(n=7) | Total<br>(n=262) |
| --- | --- | --- | --- | --- | --- | --- | --- |
| Colistin | 100% (116) | 100% (23) | 100% (100) | 100% (7) | 100% (9) | 100% (7) | 100% (262) |
| Ampicillin | 98.3% (114) | 87.0% (20) | 96% (96) | 85.7% (6) | 66.7% (6) | 100% (7) | 95.0% (249) |
| CTX | 81.0% (94) | 82.6% (19) | 61% (61) | 85.7% (6) | 66.7% (6) | 85.7% (6) | 73.3% (192) |
| Cefazolin | 84.4% (98) | 78.3% (18) | 68% (68) | 85.7% (6) | 100% (9) | 85.7% (6) | 78.2% (205) |
| Aztreonam | 77.6% (90) | 65.2% (15) | 62% (62) | 85.7% (6) | 66.7% (6) | 85.7% (6) | 70.6% (185) |
| Meropenem | 1.7% (2) | 8.7% (2) | 2% (2) | 0 | 0 | 0 | 2.3% (6) |
| Gentamicin | 60.3% (70) | 56.5% (13) | 54% (54) | 71.4% (5) | 33.3% (3) | 57.1% (4) | 56.9% (149) |
| Streptomycin | 74.1% (86) | 73.9% (17) | 53% (53) | 71.4% (5) | 22.2% (2) | 57.1% (4) | 63.7% (167) |
| Amikacin | 34.5% (40) | 8.7% (2) | 11% (11) | 57.1% (4) | 0 | 42.9% (3) | 22.9% (60) |
| Tetracycline | 95.7% (111) | 100% (23) | 98% (98) | 85.7% (6) | 33.3% (3) | 85.7% (6) | 94.3% (247) |
| SXT | 100% (116) | 100% (23) | 98% (98) | 85.7% (6) | 66.7% (6) | 100% (7) | 97.7% (256) |
| Fosfomycin | 80.2% (93) | 65.2% (15) | 61% (61) | 85.7% (6) | 11.1% (1) | 57.1% (4) | 68.7% (180) |
| Ciprofloxacin | 82.8% (96) | 82.6% (19) | 81% (81) | 85.7% (6) | 88.9% (8) | 85.7% (6) | 82.4% (216) |
| Nalidixic acid | 95.7% (111) | 87.0% (20) | 95% (95) | 85.7% (6) | 55.6% (5) | 100% (7) | 93.1% (244) |
| Chloramphenicol | 83.6% (97) | 95.7% (22) | 87% (87) | 85.7% (6) | 44.4% (4) | 100% (7) | 85.1% (223) |
| Florfenicol | 77.6% (90) | 82.6% (19) | 82% (82) | 85.7% (6) | 33.3% (3) | 28.6% (2) | 77.1% (202) |
| Nitrofurantoin | 17.2% (20) | 34.8% (8) | 19% (19) | 14.3% (1) | 0 | 0 | 18.3% (48) |

## DISCUSSION

*E. coli* is the predominant pathogen responsible for human infections [24]. To gain a more in-depth understanding of resistance genes in *E. coli*, researchers frequently undertake retrospective studies on strains preserved in laboratories [25]. This retrospective investigation determined that the prevalence of PMQR and MCR genes in *E. coli* within China was considerably high, pointing to a grave situation. Antimicrobial resistance (AMR) is exacerbated by the overuse of antibiotics, with the spread of drug resistance further intensified by numerous mobile genetic elements and their associated resistance genes.

FQ antibiotics are widely used to treat both human and animal infections, largely due to their selective inhibition of bacterial DNA synthesis. These antibiotics represent nearly one-fifth of global antibiotic consumption [26]. Our research discovered that 1,613 strains (44.0%) were PMQR-positive, with *oqxAB* being the most prevalent among PMQR genes. The detection rate of these genes in human strains was significantly lower than in other hosts. The polygenic distribution noted in our study revealed that 332 strains (9.1%) carried multiple PMQR genes concurrently, 287 (7.8%) carried two PMQR genes, and 45 (1.2%) had three PMQR genes. On the whole, the detection rate of PMQR genes in our nation was considerably lower than in many Latin American countries (for *aac(6’)-Ib-cr*: 7.3% vs. 60.6%) [27].

Since 2016, the WHO has classified colistin as one of the most critically important antibiotics. However, the global proliferation of MCR genes amongst bacteria has sparked international policy debates about the use of colistin [18,28]. In our present study, the carriage rate of MCR genes stood at 7.2%. Within this, the detection rate for *mcr-1* was 7.0%, while *mcr-3* was at 0.1%. Interestingly, one strain harbored both *mcr-1* and *mcr-3* genes. We did not identify any strains positive for *mcr-2*, *mcr-4*, or *mcr-5*. Our findings aligned with other research that pinpoints *mcr-1* as the predominant variant in Asia, succeeded by *mcr-3*. Examining host sources, the carriage rate of MCR genes in livestock surpassed that in poultry. Notably, the detection rate of *mcr-1* in pig strains peaked at 11.5%, whereas in human strains, it’s a mere 1.5%.

Considering the geographical dispersion of MCR genes, Central China exhibited a higher prevalence of *mcr-1*-positive isolates. This trend might be linked to pig farming practices and dietary preferences unique to various Chinese regions [29]. Moreover, we identified five *mcr-3*-positive isolates across three provinces: Jiangsu, Henan, and Qinghai. The absence of *mcr-2*, *mcr-4*, and *mcr-5* suggested that, despite the widespread detection of *mcr-1* in China, newer MCR gene variants have not yet permeated extensively within the country.

Considering temporal shifts, *mcr-1* was first identified in chicken strains in 2009, even though it was documented in the literature as early as 1980 [30]. Furthermore, between 2016 and 2019, five strains were found to carry *mcr-3*. Remarkably, one strain from pigs in 2017 harbored both *mcr-1* and *mcr-3*. This co-presence is an exceedingly rare occurrence in wild-type *E. coli*, with only two such strains cited in both domestic and international literature [31,32].

Our data revealed that from 2009 to 2013, the detection rate of *mcr-1* surged, peaking at 16.8% in 2013. When contrasting the periods 2007–2013 and 2000–2006, there was an 8.7-fold rise in antibiotic resistance among global *E. coli* clinical strains [33]. In response to growing concerns, the European Medicines Agency elevated the risk classification of colistin resistance from low to high in 2016. Subsequently, several nations endorsed the removal of colistin as a feed additive in livestock. This includes Brazil (November 2016), Thailand (February 2017), China (April 2017), Japan (July 2018), Malaysia (January 2019), Argentina (February 2019), and India (July 2019) [34]. Correspondingly, our study witnessed a sharp decline in the *mcr-1*-positive rate in China from 15.4% in 2016 to 5.0% by 2019. This trend underscored the pivotal role of national veterinary drug regulations in enhancing the quality of animal products, safeguarding public health, and preserving ecological safety.

Assessing the overall drug resistance of *E. coli* strains carrying MCR genes, it’s evident that all such strains exhibited multidrug resistance, with resistance rates for most drugs exceeding 50% and in some cases, surpassing 80%. Remarkably, only three drugs demonstrated a notable antibacterial effect, each with resistance rates under 30%: meropenem, amikacin, and nitrofurantoin. Of these, meropenem boasted the lowest resistance rate at just 2.3%. This finding suggested that, in clinical scenarios with multidrug-resistant Gram-negative infections, especially when accompanied by polymyxin resistance, treatment options become scarce. In such cases, carbapenems, particularly meropenem, might emerge as viable clinical alternatives.

When we considered the multidrug resistance of *E. coli* strains with MCR genes from various hosts, resistance in human strains was considerably lower than in other sources. Across categories like β-lactam, aminoglycosides, or fosfomycin, chicken strains consistently displayed higher resistance than pig strains, a disparity particularly pronounced for drugs like amikacin and fosfomycin.

All strains exhibiting MCR genes demonstrated resistance to colistin, and there was complete concordance between the MIC phenotype and genotype. Resistance levels ranged from 4 μg/mL to >64 μg/mL. This finding confirmed that the presence of MCR genes could induce colistin resistance in strains. Moreover, it underscored that the escalating detection rate of MCR in wild-type *E. coli* correlated with the rise in clinical strains resistant to polymyxin. Notably, the emergence of six strains with an MIC > 64 μg/mL pointed to the potential development of super-resistant strains. This underscored the urgency for proactive measures to address this emerging challenge.

While the rise of drug-resistant bacteria has impacted the utility of antibiotics, we cannot dispute their pivotal role in preserving both human and animal health for the foreseeable future. Monitoring antimicrobial resistance stands as a cornerstone in strategies aimed at combatting this resistance [35]. Over recent decades, China has mobilized various sectors to implement a suite of policies addressing the antibiotic resistance crisis. In August 2017, China unveiled the National Action Plan to Contain Antimicrobial Resistance (2017-2020) (http://www.moa.gov.cn/nybgb/2017/dqq/201801/t20180103_6133925.htm). Official data reflect that this holistic approach towards antimicrobial resistance has yielded significant results. Subsequently, to robustly address the issue of excessive veterinary drug residues at their source and effectively mitigate the risk posed by antibiotic-resistant animal-borne bacteria, China has rolled out a series of three-year action plans (http://www.moa.gov.cn/govpublic/xmsyj/202302/t20230228_6421716.htm). With these policies serving as a guiding force, we can anticipate improved regulation and rational application of antimicrobials across human and veterinary medicine domains. This will ensure that antibiotics are utilized judiciously in the future, further safeguarding public health.

## MATERIALS AND METHODS

### Isolation and identification of E. coli

Between 1993 and 2019, we isolated 3,663 *E. coli* strains from various sources across China. The breakdown of these isolates is as follows: humans (614), chickens (1,532), ducks (67), geese (250), pigs (846), cattle (113), goats (90), monkeys (92), rabbits (8), pigeons (25), guinea pigs (1), minks (3), cats (1), dogs (7), mice (1), squirrels (1), quails (1), river deer (2), peacocks (1), turtles (2), red pandas (2), antler deer (3), and deer (1). The samples consisted of diverse types, including meat products, milk, eggs, offal, padding, and more. These strains were collected from all 31 provinces in China, namely Beijing, Shanghai, Tianjin, Chongqing, Inner Mongolia, Xinjiang, Ningxia, Guangxi, Tibet, Heilongjiang, Jilin, Liaoning, Hebei, Henan, Shandong, Shanxi, Hunan, Hubei, Anhui, Jiangsu, Zhejiang, Fujian, Jiangxi, Guangdong, Hainan, Guizhou, Yunnan, Sichuan, Shaanxi, Qinghai, and Gansu. Identification of all strains was performed using PCR with the following primers: forward: 5’-AGAGTTTGATCCTGGCTCAG-3’; reverse: 5’-AAGGAGGTGATCCAGCC-3’. Sequencing results were then aligned using the NCBI database (http://blast.ncbi.nlm.nih.gov).

### Identification of antibiotic-resistance genes

Based on sequences from the literature and GenBank, specific primers were designed (Table S1). All isolates positive for *oqxA* were also screened for *oqxB* [20]. Two strands of the purified PCR products were sequenced, and the *qnr* alleles were identified by referencing the *qnr* gene naming conventions [21]. Additionally, all isolates testing positive for *aac(6’)-Ib-cr* were further analyzed either by FokI digestion or through direct sequencing.

### Antimicrobial susceptibility testing

In accordance with the standards recommended by the Clinical and Laboratory Standards Institute (CLSI) [22], the MICs of 17 antibiotics were determined for the 262 MCR-positive isolates using the microbroth dilution method. The quality control strain, *E. coli* ATCC 25922, was utilized to assess the efficacy of the drugs. The concentration of the antibacterial drug SXT was set at 256/4,846 μg/mL, while the concentration for other drugs was maintained at 2,048 μg/mL (Table S2).

## ACKNOWLEDGMENTS

This work was supported in part by the National Key Research and Development Program of China (2021YFD1800403), the 111 Project (D18007), and the Priority Academic Program Development of Jiangsu Higher Education Institutions (PAPD).

All authors read and approved the submitted version of the paper. We declare no conflict of interest.

## Author Contributions

Xin’an Jiao, and Xiang Chen designed the study protocol and supervised all parts of the project. Yujin Wang, Dewei Sun performed all experimental and wrote the manuscript. Zhengzhong Xu, and Xiang Chen revised the manuscript and advised the experiment. All the authors approved the final version.

